# Investigating saccade-onset locked EEG signatures of face perception during free-viewing in a naturalistic virtual environment

**DOI:** 10.1101/2024.12.12.628113

**Authors:** Debora Nolte, Vincent Schmidt, Aitana Grasso-Cladera, Peter König

## Abstract

Current research strives to investigate cognitive processes under natural conditions. Virtual reality and EEG are promising techniques combining naturalistic settings with close experimental control. However, many questions and technical challenges remain, e.g., are fixation or saccade onsets a suitable replacement as key events in continuous gaze trajectories (Amme et al., 2024), and can VR effectively capture differences across experimental conditions (Rossion & Jacques, 2008)? To address both questions, we investigate the N170 face effect in a free-viewing immersive VR study that contained houses, various background stimuli, and, notably, static and moving pedestrians to study face perception under naturalistic conditions. Our results show that aligning trials to saccade-onset leads to more well-defined ERPs, especially for the P100 component, and support that saccade-onset ERPs are the better-suited analysis method than fixation-onset ERPs for this type of experiment. Further, we observe an evolution of condition-based differences, i.e., face vs. background fixations, compatible with previous reports but extending in a large temporal window and including all electrode sites at different points in time. In summary, experiments combining VR, EEG, and eye-tracking provide further insights into the processing of faces and the relevance of saccadic onsets as event triggers under natural conditions.

**Significance Statement:** With the effort of investigating and understanding cognitive processes under naturalistic conditions, combining virtual reality and EEG can be fruitful to implement free-viewing studies. The current work combines these technologies to explore key challenges in the context of face perception in an immersive virtual environment. Our results show that saccade-onset ERPs yield more precise measurements when analyzing continuous eye-tracking data than fixation onsets. Furthermore, when processing face compared to background stimuli, distinct temporal patterns encompassing all electrode sites can be observed, offering new insights into face perception. Overall, this work highlights the potential of integrating VR and EEG to advance our understanding of cognitive processes in naturalistic settings.

## Introduction

In recent years, a step has been taken to study and understand cognitive processes under natural conditions, capturing them in dynamic, real-world environments (Gert et al., 2022; Rounds et al., 2020; Shamay-Tsoory & Mendelsohn, 2019; Stangl et al., 2023; Tromp et al., 2018). A central aspect of this approach is free-viewing paradigms, where subjects move their eyes and actively choose where to direct their gazes (Amme et al., 2024; Gert et al., 2022), allowing us to study the spontaneous and adaptive nature of real-world visual behavior (Shamay-Tsoory & Mendelsohn, 2019; Stangl et al., 2023). For these studies, virtual reality (VR) is emerging as a powerful tool, combining the high experimental control of laboratory setups with free-viewing experiences of real life (Bell et al., 2020; Bohil et al., 2011; Pan & Hamilton, 2018). Supporting the potential of VR, recent studies demonstrated that VR can provide findings similar to real life (Nolte, Hjoj, et al., 2024) and is suitable for analyzing eye-tracking data in naturalistic environments (Clay et al., 2019; Llanes-Jurado et al., 2020; Nolte, Vidal De Palol, et al., 2024). Beyond eye movements, integrating VR with electroencephalography (EEG) allows for exploring neural responses to naturalistic visual behavior (Rounds et al., 2020; Stangl et al., 2023; Tromp et al., 2018) and measuring fixation-onset event-related potentials (ERPs; Nolte, Vidal De Palol, et al., 2024). This highlights the potential of combining VR and EEG to study cognitive processes under naturalistic, free-viewing conditions.

While VR has proven helpful in investigating vision and neural processes under naturalistic conditions, many questions and technical challenges remain. For instance, although neuronal processes can be studied with VR-EEG setups (Nolte, Vidal De Palol, et al., 2024; Rounds et al., 2020; Tromp et al., 2018), the feasibility of using this combination to examine fixation-onset ERP differences across experimental conditions remains to be explored. Notably, a recent magnetoencephalography (MEG) study investigated fixation- and saccade-onset ERP differences during naturalistic viewing of pictures and found saccade-onset ERPs better suited for studying early visual components (Amme et al., 2024). Building on these findings, two questions arise: Do saccade onsets provide the optimal alignment for ERP analysis in free-viewing studies? Can VR capture differences across experimental conditions?

To address both questions, a well-established effect, such as the N170 face effect (Eimer, 2011; Rossion & Jacques, 2008), investigated in an immersive free-viewing VR study can be used. This N170 effect, described as a stronger ERP response of faces (Eimer, 2011; Rossion & Jacques, 2008) and bodies (Hietanen & Nummenmaa, 2011) compared to other stimuli, has been replicated in free-viewing picture setups (Auerbach-Asch et al., 2020; de Lissa et al., 2019; Gert et al., 2022) and with virtual humans (Wheatley et al., 2011). Thus, the N170 effect is ideal for investigating whether saccade onsets provide superior temporal alignment and whether the ERPs can reveal differences between experimental conditions in an immersive free-viewing study. The current experiment was designed as a three-dimensional virtual city populated with avatars. Participants freely explored this city center while we recorded their eye movements and EEG signals. We explored the temporal alignment of fixation- and saccade-onset ERPs in line with previous research (Amme et al., 2024) to determine the more suitable option. Furthermore, to investigate differences between experimental conditions, we split our data into head, body, and background stimuli (Gert et al., 2022). We hypothesized the highest N170 amplitude for heads, followed by bodies, and the smallest for background stimuli. Our results supported a saccade onset alignment. Noise level differences across conditions prevented directly testing the N170 effect; instead, a mass univariate analysis revealed differences between all three conditions, partially supporting our hypothesis. Overall, these results underline the suitability of combining EEG with VR but highlight new methodological challenges of free-viewing studies.

## Methods

### Subjects

Overall, 61 subjects were invited to the lab for the experiment. We could not start the recording for two subjects due to technical difficulties. Out of the remaining 59 subjects, a total of 26 subjects were excluded: five quit due to motion sickness, eighteen due to data issues during or after the recording, and finally, three subjects were excluded as they did not follow task instructions and left the central square for more than 10% of the time. This resulted in a total of 33 subjects to be included in the final dataset (19 female, zero diverse; mean age 22.63 ± 2.48). All subjects had normal or corrected to normal vision, did not report any neurological disorders, gave written informed consent before participating, and were rewarded with monetary compensation or participation hours. The ethics commission of the University of Osnabrück approved the study.

### Experimental Setup

A detailed description of the data and experimental design can be found in previous publications (Nolte, Hjoj, et al., 2024; Nolte, Vidal De Palol, et al., 2024). Below, we provide the essential aspects relevant to the current study. The experiment was developed using Unity3D (www.unity.com) version 2019.4.21f1, employing the built-in Universal Render Pipeline/Unlit with one central light source. To maintain perceptual consistency, we minimized shaded areas. The virtual environment was displayed at a constant 90 Hz frame rate via the HTC Vive Pro Eye HMD (110° field of view, resolution 1440 x 1600 pixels per eye, refresh rate 90 Hz; https://business.vive.com/us/product/vive-pro-eye-office/). Eye-tracking was facilitated using the SRanipal SDK (v1.1.0.1; https://developer-express.vive.com/resources/vive-sense/eye-and-facial-tracking-sdk/), and spatial tracking was provided by the HTC Vive Lighthouse 2.0 system (https://www.vive.com/eu/accessory/base-station2/). Participants moved within the virtual city using HTC Vive controllers 2.0 (https://www.vive.com/eu/accessory/controller2018/, sensory feedback disabled), with the direction of movement determined by the head orientation. The data was recorded on an Alienware Aurora Ryzen computer (Windows 10, 64-bit, build 19044, 6553 MB RAM; Nvidia RTX 3090 GPU, driver version 31.0.15.2698; AMD Ryzen 9 3900X 12-Core CPU). Simultaneously, EEG data using a 10/20 64-channel Ag/AgCl-electrode system with a Waveguard cap (ANT, Netherlands) and a Refa8 (TMSi, Netherlands) amplifier were recorded using the OpenVIBE acquisition server (v2.2.0; Renard et al., 2010) on a Dell Inc. Precision 5820 Tower (Windows 10, 64-bit, build 19044; Nvidia RTX 2080 Ti GPU, driver version 31.0.15.1694; Intel Xeon W-2133 CPU). The EEG data was collected at 1024 Hz with an average reference and a ground electrode under the left collarbone. Impedances were kept below ten kΩ. Synchronization between the EEG and VR systems was achieved using the LabStreamingLayer (LSL; https://github.com/sccn/labstreaminglayer). Throughout the experiment, participants were seated on a swivel chair to allow full 360° body rotation (see Figure 1A).

**Figure 1:**
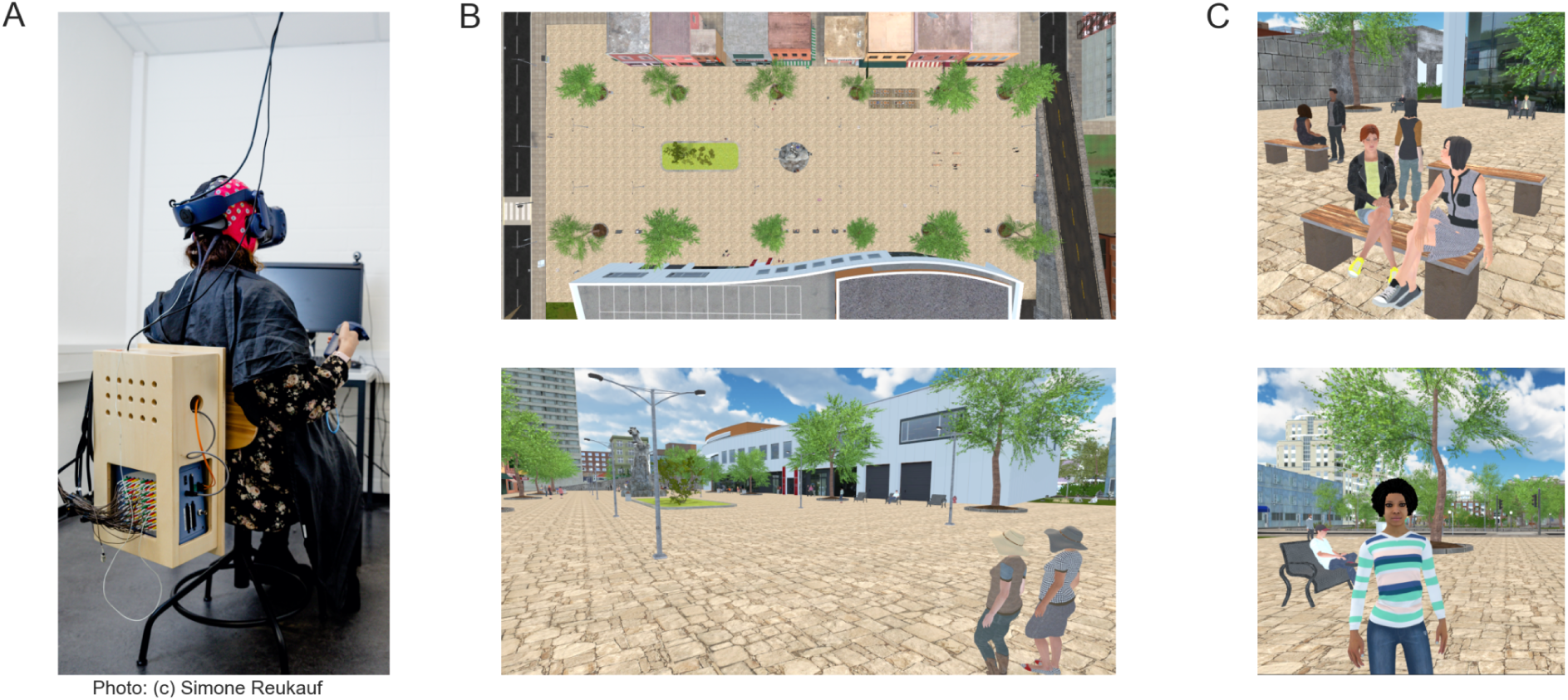
Experimental Setup. (A) Participants were seated on a swivel chair, wearing an EEG cap and VR glasses. The EEG equipment, specifically the amplifier, was stored on the back of the chair. (B) The walkable area was confined to the beige floor and comprised the center of the virtual reality scene. (C) Different pedestrians were distributed throughout the city square.

### The Experimental Procedure

The entire experiment lasted two and a half hours. At first arrival, participants filled out informed consent sheets and received instructions about the experiment. Following this, participants underwent a one-minute motion sickness test in the same virtual environment but in an unreachable part of the city. They were instructed to move toward a red sphere at the end of a street. Only participants who reported no discomfort or motion sickness after this test proceeded to the main experiment. Following this initial test, the EEG system was set up (see section “EEG Setup” for details), which took most of the time. Finally, the main experiment began with the eye tracker’s calibration and subsequent 5-point validation.

The experimental duration lasted approximately 40 minutes. The participants had 30 minutes to freely explore the central city square (see Figure 1B, marked by beige tiles) under the instruction to behave naturally as if waiting for a friend. Every 5 minutes, the exploration was paused for eye tracker validation and recalibration and for the participant to take a break if needed. After each pause, participants were returned to their previous location in the city.

### The Virtual Environment

The virtual environment was modeled to resemble a city center, populated with various background objects (e.g., buildings, foliage; see Figure 1B) and 140 pedestrians (see Figure 1C). Pedestrians were sourced from the Adobe Mixamo collection (Mixamo, 2008, https://www.mixamo.com), displaying varied activity and animation levels ranging from stationary and static to actively moving throughout the city. The pedestrians were designed to represent typical behaviors such as shopping, meeting friends, or relaxing on benches. The pedestrians did not react to the participants other than avoiding movement collisions. The active-moving pedestrians moved along predefined paths. Each object in the virtual environment had a collider, an invisible box or sphere marking the outline of an object, attached with pedestrians specifically having a separate collider for their heads and the rest of their bodies. This allowed us to separately investigate the neuronal response toward the heads and bodies of the virtual avatars. Participant movements within the virtual environment were programmed to mimic real-life displacements controlled by the participants’ head orientation and matched in speed to the moving pedestrians.

### Using gaze events to determine (EEG) trial onsets

We recorded EEG and eye-tracking simultaneously to use the timing of gaze events (fixations or saccades) as trial markers. Due to the absence of external stimulus onsets (or comparable events), we consider the data recorded during fixation (or saccade) and its immediate temporal context as a “trial”. This allowed us, for example, to investigate fixation-ERPs (Dimigen et al., 2011; Gert et al., 2022). To this end, accurate detection of event onsets (fixations and saccades) in the eye-tracking data was essential. Therefore, we employed a velocity-based eye-tracking algorithm for free viewing and free exploration in a virtual environment. This algorithm was developed to correct translational movement information superimposed on the eye movement data (Nolte, Vidal De Palol, et al., 2024; based on Dar et al., 2021; Keshava et al., 2023; Voloh et al., 2020). Applying this algorithm allowed us to differentiate between gazes (eye-stabilization movements, from now on, simply referred to as fixations) and saccades. These events could then be used as trial onsets for the EEG analysis. Specifically, we compared ERPs aligned to fixation and to saccade onset, where the latter used the saccade onset preceding the fixation as the trial onset (Amme et al., 2024). The trials for both fixation and saccade onset ERPs were split into three distinct stimulus categories: heads, corresponding to fixations on the heads of pedestrians, bodies, and background stimuli, encompassing everything that was not a pedestrian, allowing us to investigate the presence of an N170 effect in a free-viewing experiment conducted in VR. For saccade-onsets ERPs, we used the stimulus category of the fixation directly succeeding the saccades. A sequence of a participant’s walking path and a few selected fixations can be seen in Figure 2A.

**Figure 2:**
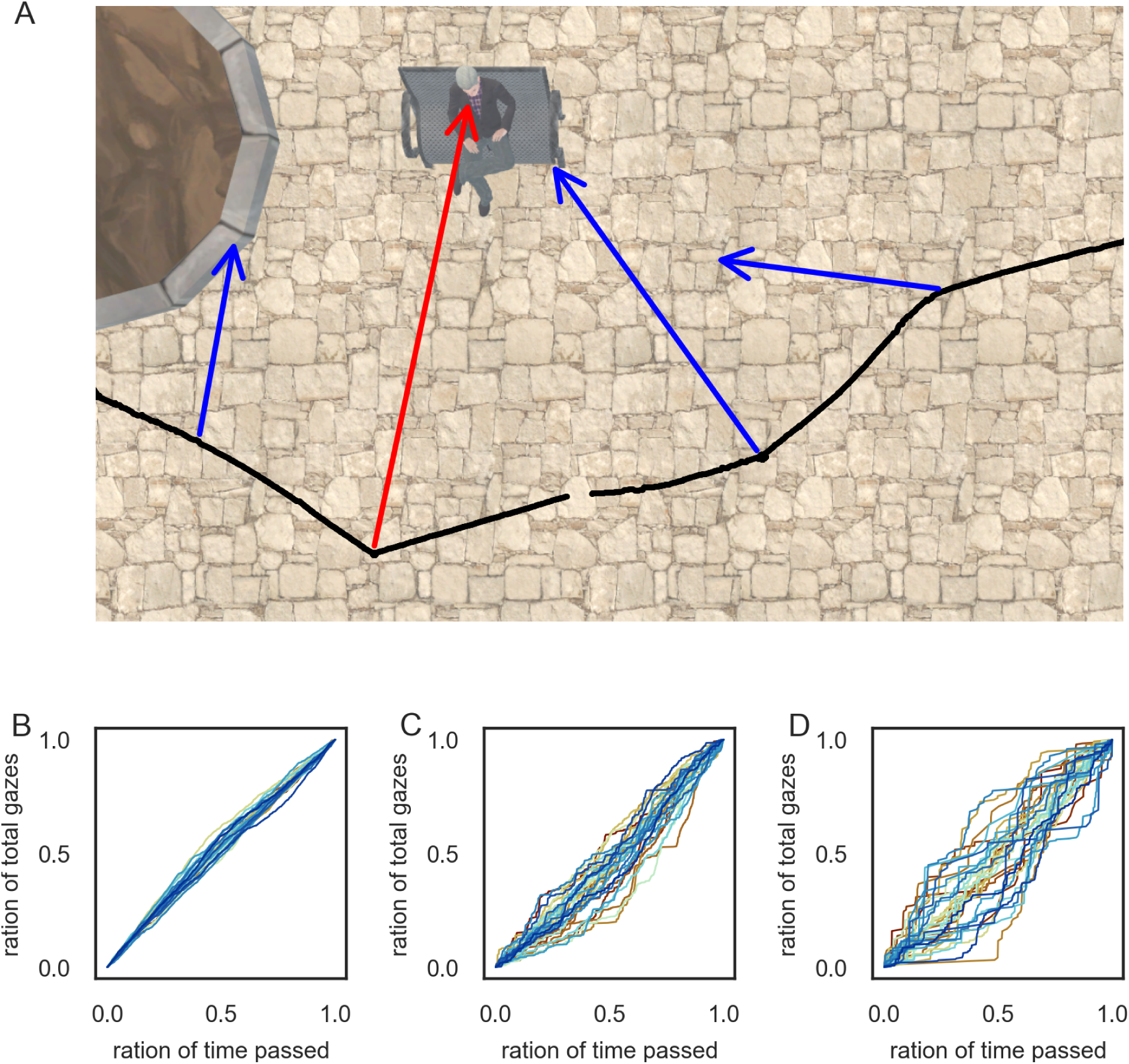
Distribution of Gaze Events. (A) An example of different fixations on different objects is plotted on top of the corresponding image of the city center. As a note, the data was slightly adjusted for visualization purposes only. The black line corresponds to the participant’s movement path, and the arrows correspond to a few selected fixations during this duration. Highlighted in red is a fixation on the face of a pedestrian. The blue arrows correspond to fixations directed at different background objects. (B)-(D) The distribution of background (B), body (C), and head (D) fixations over time, displayed as cumulative distribution functions. Each line corresponds to one participant.

To correct visible linear drifts between EEG and eye-tracking (Unity) timestamps, we adjusted the difference in timestamps between the start and end of the recordings (Nolte, Vidal De Palol, et al., 2024). As this simple adjustment did not always resolve this linear drift completely, for 20 of the subjects, the drift was additionally corrected by applying the start-end difference multiple times (with a maximum of four times) or by fine-tuning the timestamps by adding 22 ms or subtracting 11 ms milliseconds after visual inspection of the stream alignments based on fixation- and saccade-onset ERPs.

### EEG Preprocessing

Preprocessing was performed in Matlab (R2024a) using the EEGLab software (Delorme & Makeig, 2004; version 2020.0). EEG data were first loaded into Matlab, channels were renamed according to the 10-5 BESA standard system, and empty channels were removed. We then imported a separate trigger file containing all relevant fixation or saccade onset events derived from our eye tracking data (s. Determine EEG Trial Onsets for a detailed explanation). We applied a low-pass filter at 128 Hz and a high-pass filter at 0.5 Hz (pop_eegfiltnew, using a hamming window; Widmann et al., 2015). Subsequently, we downsampled the EEG data from 1024 Hz to 500 Hz. Next, we applied a line noise filter from the ‘zapline plus’ plugin (Klug & Kloosterman, 2022; based on de Cheveigné, 2020) to automatically remove spectral peaks around 50 Hz and separately 90 Hz. Then, ensuring the data was referenced to the average reference, we applied automated cleaning of noisy channels and data segments using the ‘clean_rawdata’ plugin (Kothe et al., 2019). Our data included active movement and contained more noise than expected in a classic stationary laboratory setup. To this end, we chose to apply a conservative burst criterion of 20, referring to the standard deviation cutoff for the removal of bursts via artifact subspace reconstruction (ASR). Removed noisy segments were saved to be used by the unfold toolbox (Ehinger & Dimigen, 2019; for a detailed description, see EEG Analysis). After channel removal, the clean dataset was re-referenced to the average reference once more. Using the AMICA plugin (version 15, Palmer et al., 2012), we performed an independent component analysis (ICA) on the cleaned data to identify and remove muscle, eye, heart, or remaining line or channel noise. For this step only, we high-pass filtered our data to 2 Hz (Dimigen, 2020). Components containing more than 80% muscle activity or more than 90% of other noise, as identified by ICLabel (Pion-Tonachini et al., 2019), were removed automatically. Finally, we interpolated the missing channels (spherical interpolation). The described procedure was repeated for all subjects before we applied further statistical analysis.

### EEG Analysis

First, we analyzed and compared ERPs aligned to fixation and saccade onsets, investigating the ERP waveforms for -300 to 500 ms surrounding each event for individual subjects and the averaged ERPs across subjects.

Next, to account for and correct the effect of overlapping events due to our free-viewing paradigm, we used a linear model implemented by the unfold toolbox (Ehinger & Dimigen, 2019) with the current event factor and the levels of background, body, and head. This overlap correction was applied for -500 up to 1000 ms surrounding saccade onsets (Gert et al., 2022).

To investigate differences across conditions at all electrodes and time points (-500 to 1000 ms surrounding saccade onset), we conducted a one-factor repeated measures ANOVA (1x3: Head, Body, and Background), with an alpha level set at .05. To account for the multiple comparison problem, we applied a cluster-based permutation test incorporating threshold-free cluster enhancement (TFCE), as implemented via the ept_TFCE Matlab toolbox (Mensen & Khatami, 2013). We performed 10,000 permutations, randomizing data across the three factors for each permutation, followed by a one-factor repeated measures ANOVA. The resulting F-values were enhanced using TFCE (parameters E = .666, H = 1) based on recommendations for F-statistics (Mensen & Khatami, 2013). This process generated an empirical null distribution (H0) of TFCE-enhanced F-values, and the maximum F-value across channels and time points for each permutation was recorded. The observed TFCE-enhanced F-values were then compared to the empirical distribution, with statistical significance determined as values exceeding the 95th percentile of the null distribution.

## Results

### Gaze Events

Before analyzing ERPs, we first compared the different gaze events by examining the median and median absolute deviation (MAD) across various aspects of each category. One clear difference between the three categories - background stimuli, bodies, and heads - was the number of trials. Background stimuli had the most trials, with a median of 3,968 ± 492.224. Notably, the body and head categories had approximately ten times fewer trials than the background condition, with bodies averaging 741 ± 220.908 trials and heads 151 ± 133.434 trials. Although there was considerable between-subject variation in the number of trials for both the body (min = 290, max = 1296) and head (min = 17, max = 913) categories, the number of fixations directed at bodies and heads was not significantly correlated across participants (r = .184, p = .305). This indicated that it was not simply that some subjects gazed at pedestrians overall more or less; instead, some subjects focused more on heads while others directed more fixations toward bodies. Despite the differences in trial counts, fixations in all three conditions were equally distributed across the entire experimental duration (see cumulative distribution functions, Figure 2B-D). This balanced distribution was essential, enabling comparison across the three conditions without adjusting for differences in experimental duration or participant fatigue. When examining the median and MAD for event durations, fixation durations were similar across all three categories: background stimuli (.186 ± .013 s), bodies (.2 ± .016 s), and heads (.178 ± .023 s). In contrast, saccade durations differed, with background stimuli having the longest saccades (.076 ± .003 s), followed by bodies (.066 ± .001 s), and heads with the shortest saccades (.056 ± .015 s). A similar pattern emerged for saccade amplitudes, where saccades towards background stimuli had the highest amplitudes (12.119 ± 2.104), followed by bodies (7.212 ± 1.618) and heads (5.35 ± 1.491). These findings highlighted that, despite similarities in fixation durations and their temporal distribution, the different stimulus categories were associated with different saccade patterns and varying numbers of events.

### Comparing fixation- and saccade-onset ERPs

In investigating ERPs, we first examined a single subject to compare the difference between fixation and saccade onset ERPs, which aligned with Amme and colleagues (2024). For this purpose, we selected a representative subject, with a total of 4620 trials, split into 3779 background, 747 body, and 194 head trials. To compare the difference between fixation- and saccade-onset ERPs, we sorted the subject’s fixation-onset trials according to saccade durations (Figure 3A; Amme et al., 2024). The curved amplitude peaks highlighted that the P100 component followed saccade rather than fixation onsets, promoting saccade onsets as the more suitable time points for aligning individual trials in our free-viewing experiment. This notion was further supported by more temporally focused saccade-onset ERPs with time-shifted but higher P100 amplitudes than their smaller, more smeared-out fixation-onset ERP counterparts (Figure 3B). Notably, for this subject, the saccade-onset ERP waveform of the head condition was visibly different from the other two conditions. First, we observed a different level of noise for this condition, most likely due to the smaller number of trials. Secondly, the P100 was smaller for the head than the other two conditions. However, comparing the P100 amplitudes across subjects highlighted the similarity between the background and body conditions (Figure 3C) but also between background and head (Figure 3D) and body and head (Figure 3E) conditions, demonstrating that the difference in the P100 amplitude for this subject was not a pattern observable across subjects. Overall, the single-subject results supported the idea that saccade-onset ERPs might be the better-suited analysis method than fixation-onset ERPs for this type of experiment.

**Figure 3:**
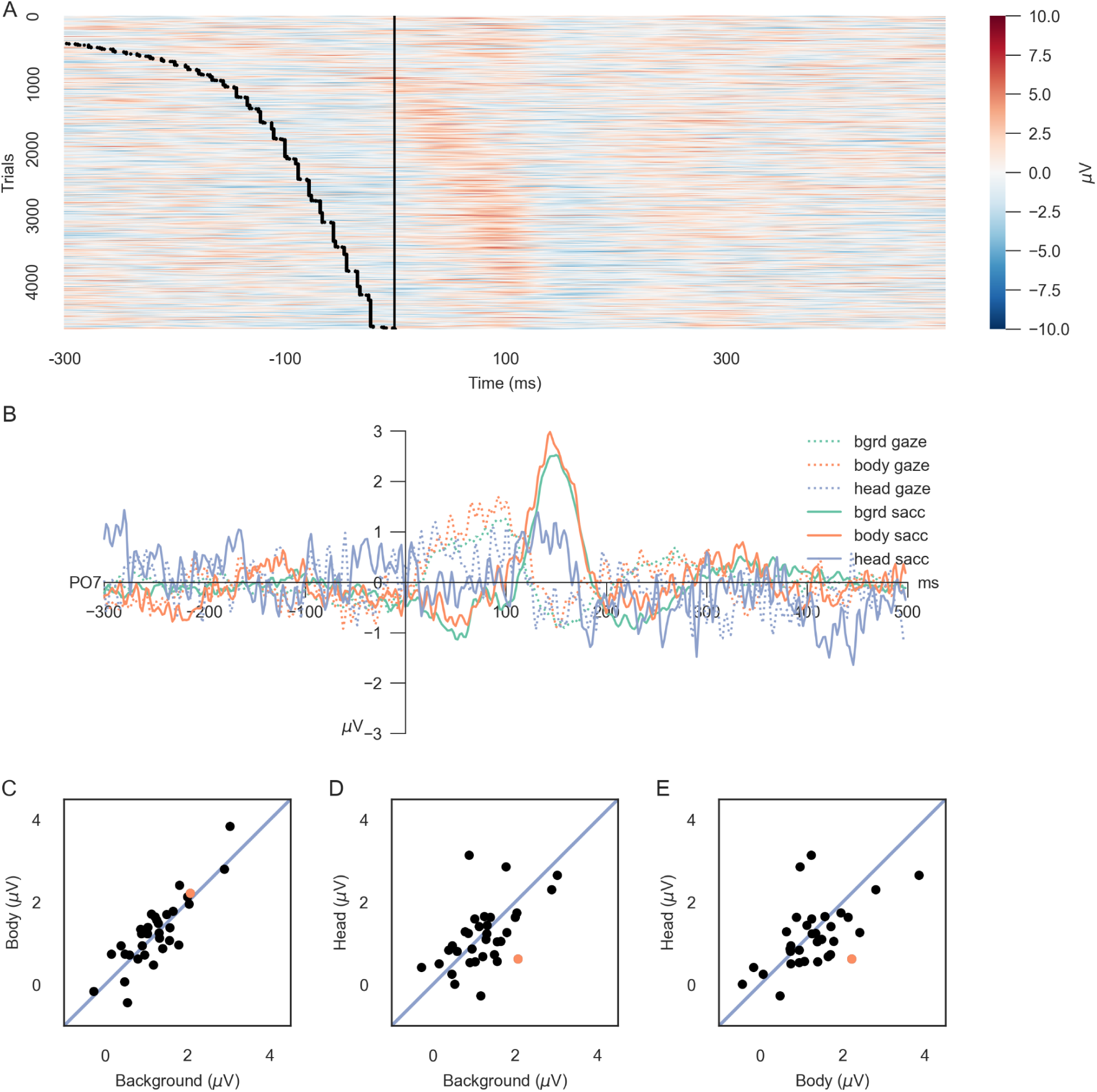
Fixation and saccade onset ERPs for a single subject. (A) All trials of one subject at electrode PO7, aligned to fixation onset, are sorted according to saccade duration, with the first trials (top of the y-axis) having the longest saccade durations. Red indicates positive, blue negative amplitudes. The trials are plotted over time. The black vertical line corresponds to fixation and, therefore, trial onset. The trials are smoothed for visualization with a Gaussian filter of 2 in the y and a Gaussian filter of 5 in the x direction. The black dotted line represents the saccade onset in each trial. (B) Fixation- (dotted lines) and saccade-onset (solid lines) ERPs of the same subject for electrode PO7. The different categories, background, body, and head, are indicated by the different colors. (C) - (E) For the saccade-onset ERPs, the mean amplitude between 150 - 170 ms of each subject’s average ERP is plotted for (C) background vs. body, (D) background vs. head, and (E) body vs. head trials. Each dot represents one subject. The single subject used in (A) and (B) is highlighted in orange. The blue lines in the background indicate a (potentially) perfect linear relation between the two conditions.

Next, we investigated the difference between fixation- and saccade-onset ERPs across subjects. Fixation-onset ERPs (dotted lines, Figure 4A) show a broad P100 component across all three stimulus categories, with background stimuli evoking the highest P100 peak and head stimuli the lowest. In comparison, saccade-onset ERPs (solid lines, Figure 4A) are shifted in time but exhibit higher amplitudes across all stimulus categories and P100 peaks that are more temporally focused. Interestingly, the differences in P100 peaks between the stimulus categories seen in fixation-onset ERPs disappear with saccade-onset alignment. Like the single subject results, the head condition appears to be the noisiest, regardless of the alignment. The topographical analysis across all channels (Figure 4B & C) supported this observation, highlighting that the preference for saccade-onset ERPs is not restricted to only the single, selected electrode. ERPs aligned to saccade onset elicit higher amplitudes across occipital electrodes than those aligned to fixation onset. These findings support the usage of saccade-onset versus fixation-onset alignment in EEG analyses due to their impact on the overall ERP curve. Saccade-onsets lead to more well-defined ERPs, especially concerning the P100 component.

**Figure 4:**
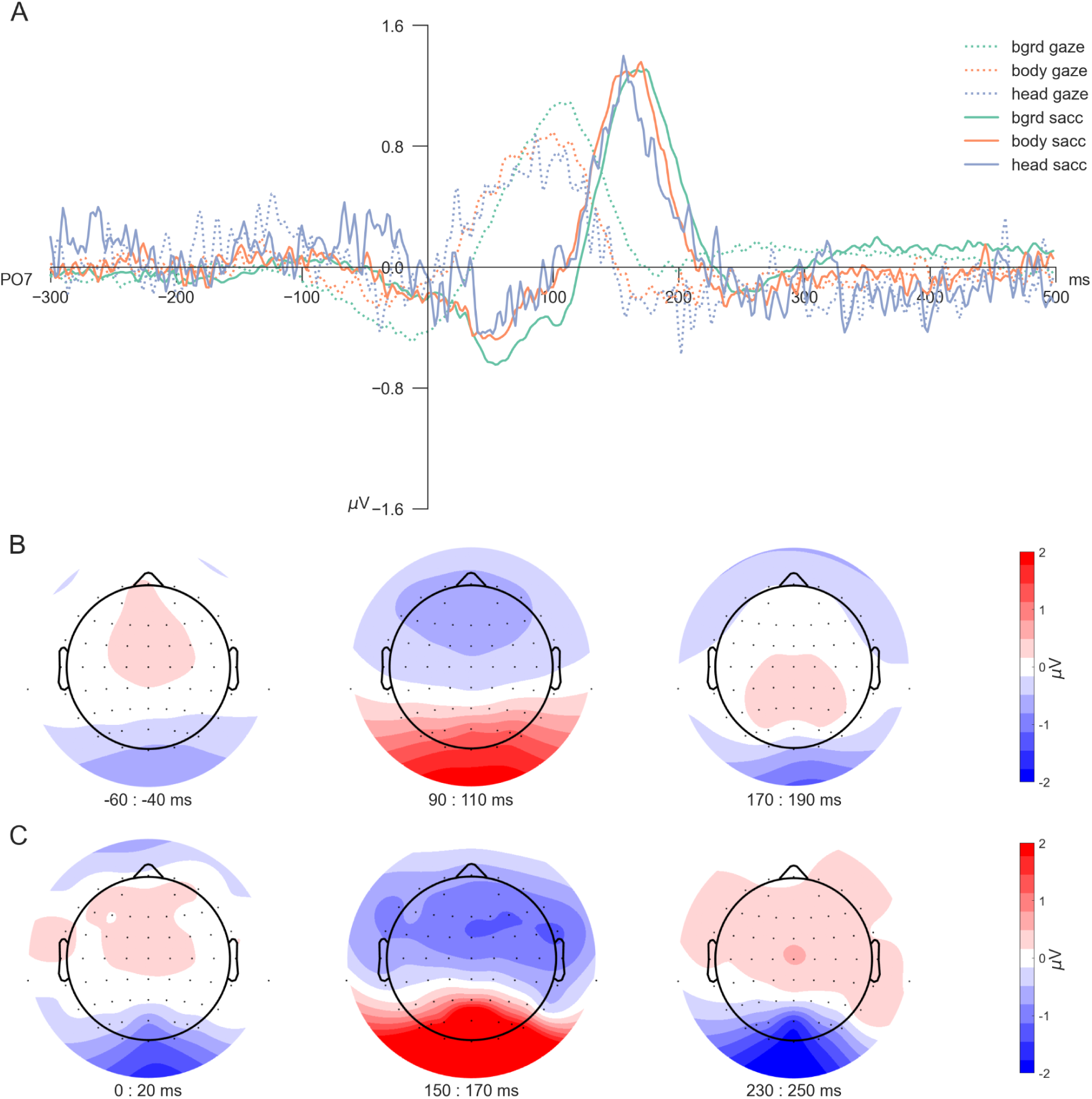
Fixation vs saccade onset ERPs. (A) Across subject ERPs at channel PO7 for all three conditions for the two different onsets: fixation-onset ERPs shown with dotted lines, saccade-onset ERPs with solid ones. (B) and (C) Topoplots of (B) fixation-onset and (C) saccade-onset ERPs, shown for three distinct time points: before the P100, around the P100, and at the N170.

### Comparing head, body, and background trials

To examine condition-specific differences, we first focused on channels discussed in previous literature (e.g., Gert et al., 2022), particularly PO7 (Figure 5A) and PO8 (Figure 5B).

**Figure 5:**
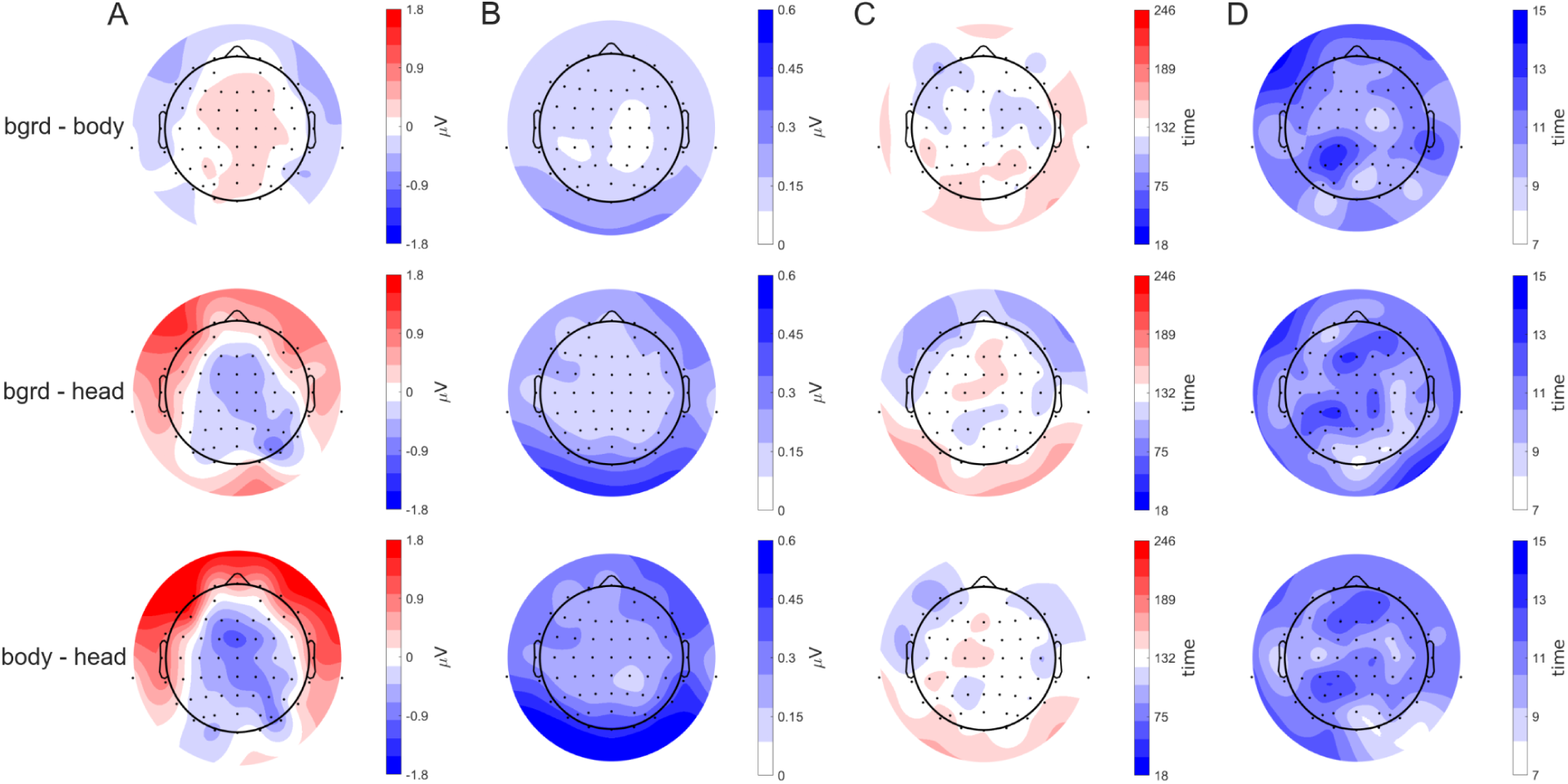
ERPs for all three conditions. (A)-(D) The across-subject average saccade-onset and deconvoluted ERPs were separated for background, body, and head trials. The average ERPs and corresponding confidence intervals are shown for four different electrodes: PO7 (A), PO8 (B), P2 (C), and F7 (D). The conditions are indicated by different colors. (E) - (G) Difference plots of two conditions at distinct time points after saccade onset: 20 ms, 106 ms, 150 ms, 200 ms, and 240 ms. The topographic plots are shown for the average difference between (E) background minus body, (F) background minus head, and (G) body minus head trials.

Comparing the deconvoluted potentials across conditions at these channels revealed no visible discrepancies, with only minimal, if any, negative deflection following the initial positive peak, a time-shifted P100. In contrast, other electrodes displayed notable differences between conditions. For instance, at electrode P2 (Figure 5C), the head trace diverged from the body and background traces after saccade onset until reaching peak amplitudes around 150 ms. Similarly, at frontal sites such as F7 (Figure 5D), distinctions between the head compared to both background and body conditions were visible around fixation onset, approximately 50-80 ms after saccade onset. Notably, across all four electrodes, the head condition exhibited a higher level of noise and more considerable between-subject variability than the other two conditions, most likely caused by the lower number of trials. This variability of the head condition and the observed topographical distinctions required a statistical approach beyond traditional measures such as peak-to-peak comparisons.

Accordingly, we employed a mass univariate analysis to examine condition-specific differences across electrodes and time. Specifically, testing for significant differences using TFCE (correction for multiple comparisons with α < .05) revealed a significant cluster most compatible with an effect spanning from 18 ms to 246 ms and encompassing all electrodes. The cluster started at channel F7 at 106 ms (see Figure 5D and the second difference plot in Figures 5E-G) shortly after fixation onsets, with a median difference between background and body of -.174 μV, a median difference between background and head of .505 μV, and between body and head of .658 μV at this time point and electrode. Inspecting temporal differences between conditions further highlighted this effect (Figure 5E-G), revealing a gradual shift in condition differences, primarily driven by the head condition (Figure 6A). Notable, while the maximum contribution is primarily due to differences between the head and either of the other two conditions (Figure 6A and 6B), the timing of each maximum contribution differs little for all three contrasts. Specifically, it can be observed that frontal electrodes have their highest contribution on average around saccade onset, while occipital electrodes have their highest contribution slightly later (Figure 6C and 6D). Overall, these findings support the presence of condition-based differences throughout the entire temporal window and all electrode sites, with the highest differences visible around saccade onset.

**Figure 6:**
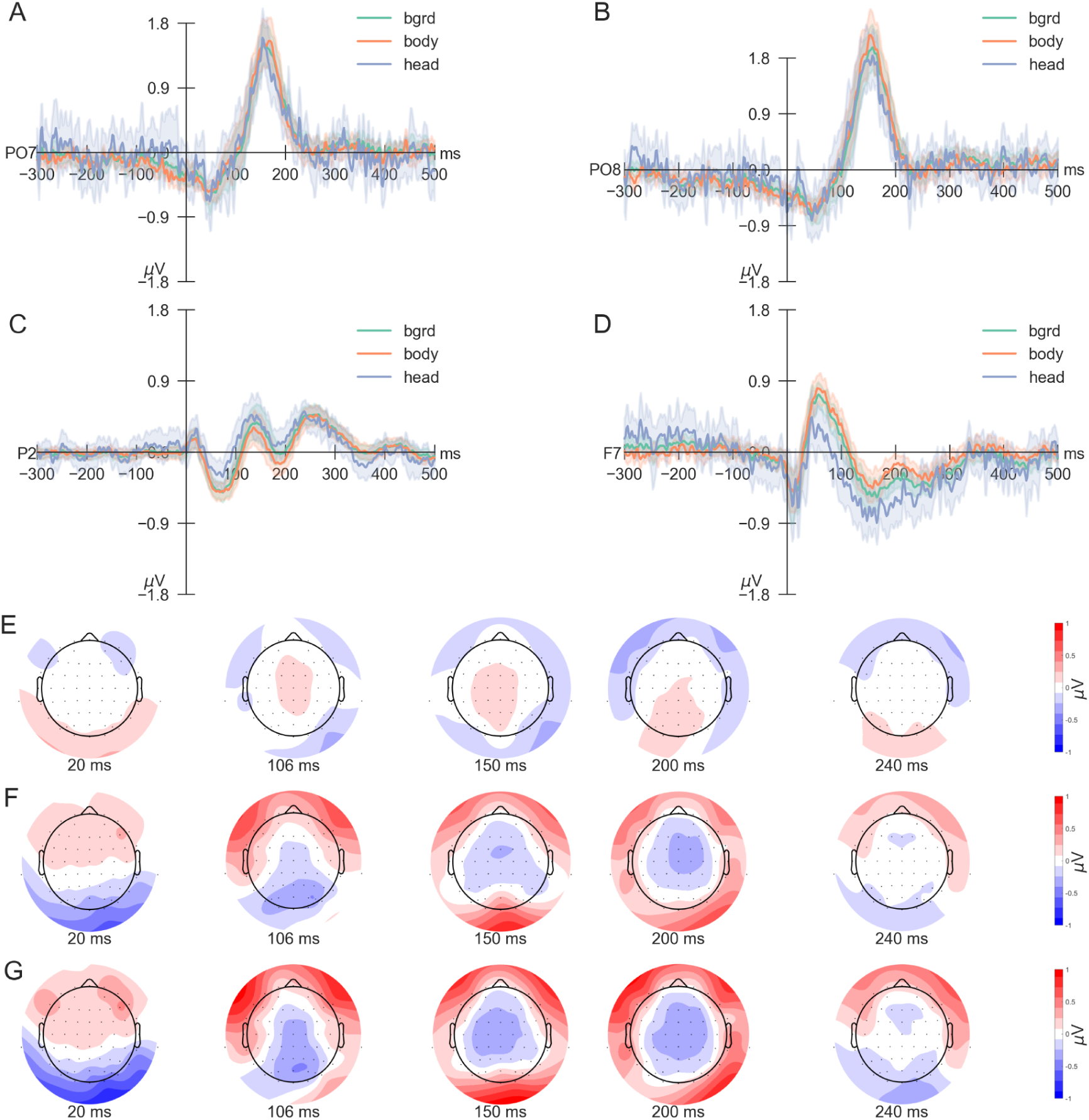
Measures of effect size. For each of the three differences (background-body, background-head, body-head), the results of the TFCE are displayed. (A) The maximum contribution averaged over participants is displayed for each electrode. The results are given in μV. (B) The corresponding standard errors of the mean for each maximum distribution are displayed. (C) The time points for the maximum contributions of each electrode are shown. The color bar corresponds to the time interval of the significant cluster, lasting from 18 to 246 ms. (D) The corresponding standard error of the mean for the time of each maximum contribution is displayed.

## Discussion

This study explored methodological considerations for using virtual reality and EEG to investigate event-related potentials in a free-viewing task. We designed a three-dimensional virtual city populated with virtual pedestrians, allowing participants to navigate and visually explore as they would in a real-world setting. Our findings align with a recent MEG study (Amme et al., 2024), showing that saccade-onset locked ERPs offered more consistent and interpretable results than traditional fixation-onset ERPs, particularly during time windows associated with P100 responses. These results support the critical role of self-initiated eye movements in priming the brain for incoming visual information (Amme et al., 2024) and advocate for a methodological shift in free-viewing paradigms toward saccade-onset ERPs to improve our understanding of visual processing.

We analyzed differences across experimental conditions by calculating saccade-locked ERPs using eye movements generated and recorded in virtual reality. Specifically, we aimed to replicate the well-established N170 face effect (Eimer, 2011; Rossion & Jacques, 2008). Our results partially supported our hypothesis, showing a significant cluster of differences among the three experimental conditions: heads, bodies, and background stimuli. These differences spanned much of the trial duration, including the P100 and N170 time windows across all electrodes, consistent with previous research (Gert et al., 2022) and therefore supporting the feasibility of VR for examining condition-based differences. Although we did not model saccade amplitudes, which could have contributed to the observed differences between conditions (Ehinger & Dimigen, 2019), the extended temporal cluster beyond the P100 component implies that neuronal processes associated with face processing (Freiwald et al., 2016; Gao et al., 2019; Rossion & Jacques, 2008) likely contribute to these differences. These results demonstrate the feasibility of investigating differences between experimental conditions in VR using free-viewing paradigms.

Notably, while we observed differences across experimental conditions, directly testing our hypothesis regarding the N170 effect using standard methods, such as the peak-to-peak effect (i.e., Gert et al., 2022), was impossible. This was caused by varying noise levels across the three conditions, most likely due to differences in trial counts. While viewing faces is known to lead to substantial inter-individual differences in gaze behavior (Guy et al., 2019; Guy & Pertzov, 2023; Rubo et al., 2023), the high between-subject variability and a low number of head fixations for some subjects were surprising, especially since the pedestrians were the only stimuli displaying changes in movement, something known to capture attention (Abrams & Christ, 2003). This underscores a fundamental challenge for free-viewing studies: the inability to control trial counts per condition or ensure what will be seen by the subjects. Future studies could increase the experimental duration, though restricted by experiences of motion sickness and participant fatigue (Clay et al., 2019; Nolte, Vidal De Palol, et al., 2024), or increase the frequency of relevant stimuli to ensure a satisfying number of trials per condition. It remains uncertain, however, whether increasing the trial count in the current study would lead to conclusive results comparable to previous research (Auerbach-Asch et al., 2020; Gert et al., 2022). Visible inspection of our ERP traces revealed an absence of negative peaks in occipital-temporal electrodes typically implicated in N170 research (Eimer, 2011; Gert et al., 2022; Rossion & Jacques, 2008). Several reasons come to mind. First, N170 effects might be more individual, with hemispheric differences affecting ERP responses (De Vos et al., 2012), rendering an across-subject investigation difficult. Second, free-viewing studies often show smaller ERP components than stimulus-locked studies (Auerbach-Asch et al., 2020; Gert et al., 2022; Ladouce et al., 2022), a difference we may also observe in VR-based free-viewing studies. Finally, we did not control for directional effects of heads, which are known to influence N170 amplitudes (Eimer, 2000; Rossion & Jacques, 2008), as limited head-fixation data constrained our ability to conduct a robust analysis. These results highlight challenges when adapting lab-based methodologies for free-viewing studies and suggest the need for a methodological shift when moving towards free-viewing experiments.

In conclusion, our results demonstrated the potential of VR-based, free-viewing paradigms combined with EEG to capture meaningful neural data, with saccade-onset-locked ERPs offering advantages over fixation-onset for ERP analysis. Employing saccade-onset ERPs, we observed significant differences between experimental conditions, providing a suitable approach for analyzing free-viewing data. However, variability in noise levels across conditions and among subjects underscores the complex nature of free-viewing studies. While combining free-viewing in VR with EEG recordings is feasible and yields valuable data, significant methodological challenges remain. With this work, we highlight the need to develop further and refine measurement and analysis approaches to suit the needs of advanced experimental designs. Thereby, we aim to contribute to laying a foundation for future studies to advance the field of free viewing or exploration paradigms in visual 3D environments further.

## Acknowledgements

The authors would like to thank everyone who contributed to the project. Specifically, they would like to thank John Madrid-Carvajal, Jakob Litsch, Eva von Butler, Anneke Büürma, Marketa Becevova, Reem Hjoj, and Marie Bensien for help in collecting the data.

Furthermore, the authors would like to thank Marc Vidal De Palol, Jakob Litsch, and Anna L. Gert for their support in designing the study. The authors would like to thank Artur Czeszumski for his input while developing the automated EEG preprocessing pipeline and Jessica Simon for investigating different preprocessing parameters. The authors would also like to thank Moritz Lönker for implementing the saccade amplitude calculations. Finally, the authors would like to thank Tracy Sánchez Pacheco for her input and feedback on the visualizations and TFCE analysis.

